# JAK1 palmitoylation by ZDHHC3/7 is Essential for Neuropoietic Cytokine Signaling and DRG Neuron Survival

**DOI:** 10.1101/2020.11.16.385971

**Authors:** Luiselys M. Hernandez, Audrey Montersino, Jingwen Niu, Shuchi Guo, Gareth M. Thomas

**Affiliations:** Shriners Hospitals Pediatric Research Center (Center for Neurorehabilitation and Neural Repair), Lewis Katz School of Medicine at Temple University, 3500 N. Broad Street, Philadelphia, PA 19140; Department of Anatomy and Cell Biology, Lewis Katz School of Medicine at Temple University, 3500 N. Broad Street, Philadelphia, PA 19140

## Abstract

Janus Kinase-1 (JAK1) plays key roles in pro-survival signaling during neurodevelopment and in responses to neuronal injury. JAK1 was identified as a potential palmitoyl-protein in high-throughput studies, but the importance of palmitoylation for JAK1’s roles in neurons has not been addressed. Here, we report that JAK1 is endogenously palmitoylated in cultured Dorsal Root Ganglion (DRG) neurons and, using an shRNA knockdown/rescue approach, reveal that JAK1 palmitoylation is important for neuropoietic cytokine-dependent signaling and neuronal survival. We further identify the related palmitoyl acyltransferases (PATs) ZDHHC3 and ZDHHC7 as dominant regulators of JAK1 palmitoylation in transfected non-neuronal cells and endogenously in neurons. At the molecular level, palmitoylation is critical for JAK1’s kinase activity in transfected cells and even *in vitro,* likely because palmitoylation facilitates transphosphorylation of key sites in the JAK1 activation loop. These findings provide new insights into palmitoylation-dependent control of neuronal development and survival.

## Introduction

During neurodevelopment, the axons of Dorsal Root Ganglion (DRG) neurons extend into peripheral tissues where limiting amounts of trophic factors stimulate axon-to-soma signals that support neuronal survival. The best-known example of this process is the Nerve Growth Factor (NGF) -dependent engagement of the tyrosine kinase TrkA on the surface of axons (Harrington and Ginty 2013). However, there is evidence that additional signals are also important for DRG neuron survival (Nakashima et al. 1999; Rodig et al. 1998). These include pathways activated by neuropoietic cytokines, including Interleukin-6 (IL-6), Ciliary Neurotrophic Factor (CNTF), Leukemia Inhibitory Factor (LIF), Cardiotrophin-1 (CT-1) and Oncostatin-M (OSM) (Bauer et al. 2007). IL-6 family cytokines bind receptor complexes containing the transmembrane protein Gp130 (gene name *IL6ST),* triggering autophosphorylation of Janus Kinases (JAKs) and subsequent phosphorylation of ‘downstream’ JAK substrates, most notably the Signal Transducer and Activator of Transcription (STAT) family of transcription factors (Bauer et al. 2007). There is evidence that gp130-JAK-STAT signals act in a similar retrograde manner to those characterized for NGF/TrkA (Hendry et al. 1992), and JAK-STAT retrograde signaling can be directly demonstrated in peripheral neurons in culture (Bareyre et al. 2011; Ben-Yaakov et al. 2012; O’Brien and Nathanson 2007). Moreover, Gp130/JAK signaling in DRG neurons is implicated not only during neurodevelopment, but also in responses of the developed nervous system to axonal injury, and in pathological conditions such as chronic itch (Lu et al. 2014; Oetjen et al. 2017; Rodig et al. 1998). However, in contrast to the wealth of knowledge regarding NGF/TrkA signaling, remarkably little is known regarding how neuropoietic cytokine/JAK signaling is controlled in neurons.

The importance of Gp130/JAK signaling for DRG neuron survival is exemplified by the perinatal death and reduced number of DRG neurons seen in mice lacking either *Jak1* or *Gp130* (Nakashima et al. 1999; Rodig et al. 1998). Importantly, DRG neurons from each of these mouse lines survive poorly in culture when neuropoietic cytokines are used as a source of trophic support (Nakashima et al. 1999; Rodig et al. 1998), suggesting that *in vivo* DRG neuronal loss in intact *Jak1* or *Gp130* knockout (KO) mice is cell autonomous. It is, however, unclear why other JAK family members (JAK2, JAK3 and the related TYK2 (Babon et al. 2014; Leonard 2001)) cannot compensate for loss of *Jak1,* particularly given their broadly similar domain structure and substrate specificity (Briscoe et al. 1996; Harpur et al. 1992; Manning et al. 2002). We thus hypothesized that JAK1 may be subject to differential regulation, perhaps by different post-translational modification, compared to other JAK family members.

In this regard, we were struck by numerous proteomic studies that suggested that JAK1 is subject to palmitoylation, the covalent modification with the lipid palmitate (Collins et al. 2017; Morrison et al. 2015; Ren et al. 2013; Sobocinska et al. 2018; Thinon et al. 2018; Wei et al. 2014). Palmitoylation is best known to target proteins to specific membranes, but can also impact additional aspects of protein interactions and, in the case of kinases, enzymatic activity (George et al. 2015; Holland et al. 2016; Linder and Deschenes 2007; Resh 2006; Runkle et al. 2016). In contrast to the frequent reports of likely JAK1 palmitoylation, JAK2 and JAK3 are absent from palmitoyl-proteomic databases and Tyk2 was identified in only a single palmitoyl-proteomic study (Blanc et al. 2015; Sanders et al. 2015; Thinon et al. 2018). Despite this striking difference from other JAK family members, follow-up studies of JAK1 palmitoylation have been limited to an examination of distribution of transfected JAK1 in heterologous cells (Ren et al. 2013). The potential palmitoylation of JAK1 in neuronal cells and its functional roles in the nervous system thus remain unexplored, and whether and how palmitoylation impacts JAK1’s signaling ability has likewise remained unaddressed. In prior work, we reported a surprising finding that palmitoylation is essential for the enzymatic activity, even *in vitro* of Dual Leucine-zipper Kinase (DLK) (Holland et al. 2016). The lack of available reagents, such as phosphospecific antibodies, to probe DLK’s phosphorylation and activation state prevented us from better defining how palmitoylation regulates this kinase. However, we were aware that an array of well characterized phospho-antibodies against JAKs might provide further insight into palmitoylation-dependent control of JAK activity, should we observe it.

Here we report that JAK1 is robustly palmitoylated in embryonic DRG neurons and that this modification is critical for LIF-induced DRG neuron signaling and survival. We identify the closely related palmitoyl acyltransferases (PATs) ZDHHC3 and ZDHHC7 as dominant regulators of JAK1 palmitoylation. At the molecular level, we reveal a critical role for palmitoylation in the control of JAK1 kinase activity, even *in vitro.* Consistent with this requirement, we provide evidence that palmitoylation is critical to permit transphosphorylation at key sites in JAK1’s activation loop. These findings reveal unexpected roles for palmitoylation in the control of JAK/STAT signaling and DRG neuron survival.

## Results

As a first step to determining potential roles of JAK1 palmitoylation, we assessed whether JAK1 is indeed palmitoylated. We therefore expressed HA-tagged wild type JAK1 (HA-JAK1WT) in HEK293T cells and performed a non-radioactive palmitoylation assay, Acyl Biotin Exchange (ABE; (Thomas et al. 2012; Wan et al. 2007)). We detected a strong signal for palmitoyl-HA-JAK1WT, which was absent when cells were pre-treated with a broad spectrum palmitoylation inhibitor, 2-bromopalmitate (2Br; (Jennings et al. 2009)), or when ABE samples were prepared in the absence of the key reagent hydroxylamine (NH_2_OH)(Fig 1A, B). The palmitoyl-JAK1 signal was also abolished when two cysteines that lie in the linker region upstream of JAK1’s pseudokinase domain were mutated to non-palmitoylatable serine (Cys541, Cys542, ‘HA-JAK1-CCSS’) (Fig 1A, B), consistent with a prior report (Ren et al. 2013). Together, these findings suggest that JAK1 is indeed palmitoylated.

**Figure 1:**
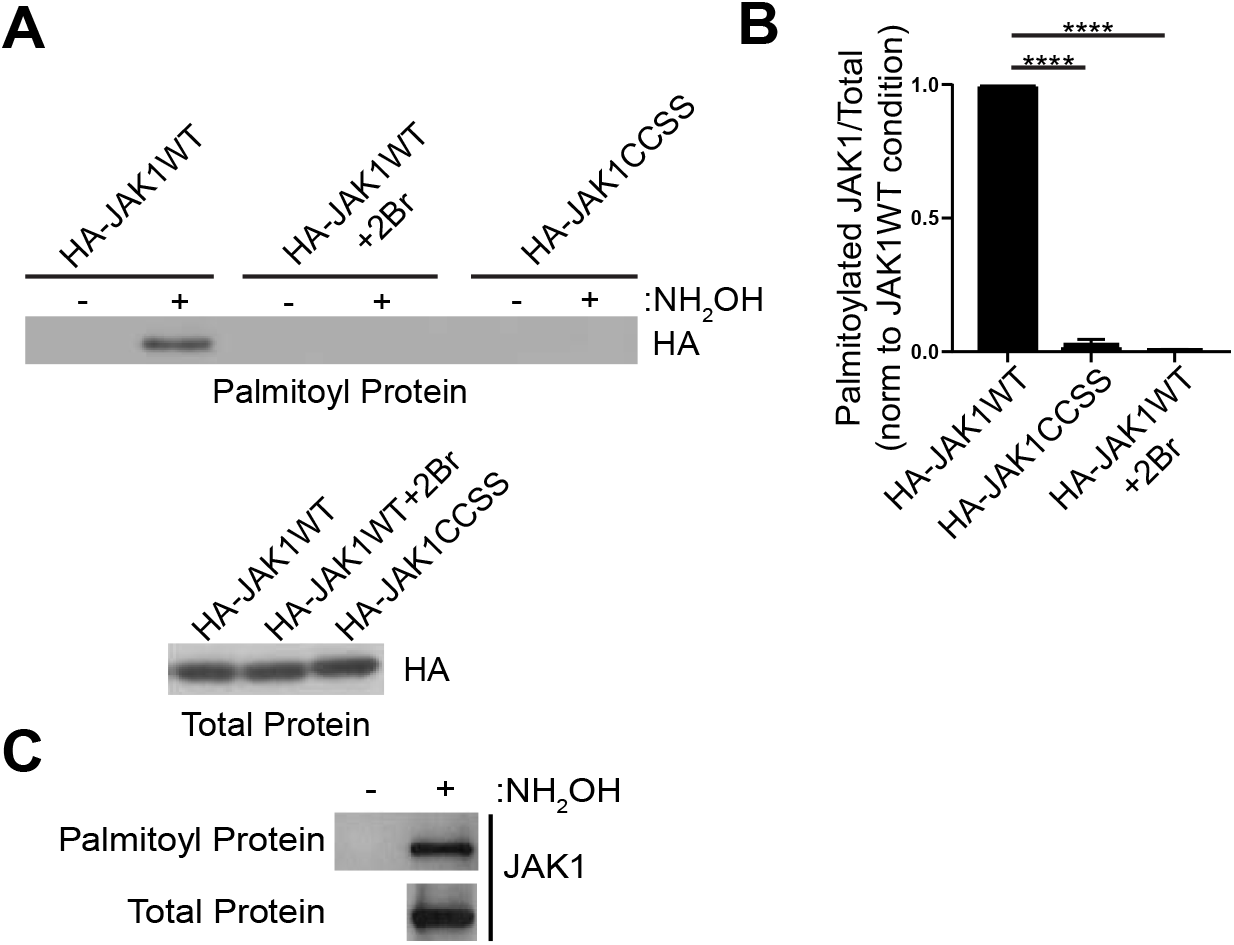
JAK1 is palmitoylated in transfected HEK293T cells and in DRG neurons. ***A***: Anti-HA western blots of palmitoyl-fractions (isolated by ABE, upper panel) and total lysates of HEK293T cells transfected with the indicated JAK1 cDNA constructs and treated with or without 2-Bromopalmitate (2-Br). ***B:*** Quantified data from *A* confirm robust palmitoylation of JAK1WT that is prevented by 2-Br treatment and by CCSS mutation. N=3-7 per condition. ****: p<0.0001 versus HA-JAK1WT, one-way ANOVA with Bonferroni *post hoc* test. All data in this and subsequent Figuregs are mean +SEM.

### Palmitoylation is important for JAKI-dependent signaling and survival of DRG neurons

We next asked whether endogenous JAK1 is palmitoylated in cultured Dorsal Root Ganglion (DRG) neurons. We observed a strong signal for palmitoyl-JAK1 in ABE fractions from cultured DRG neurons, which was absent in ABE samples prepared in the absence of hydroxylamine (Fig 1C). These findings suggest that JAK1 is endogenously palmitoylated in DRG neurons.

To determine whether JAK1 is important for DRG neuron survival, we identified an shRNA that potently reduces JAK1 protein expression. We found that lentiviral infection with this shRNA, but not a control scrambled shRNA, markedly reduced survival of LIF-supported DRG neuron cultures (Fig 2A, B, Fig S1A). This finding is reminiscent of the impaired survival of DRG neurons cultured from *Jak1* KO mice in the presence of LIF (Rodig et al. 1998) and further suggests that this impaired survival of DRG neurons cultured with LIF is due to a cell autonomous requirement for JAK1.

**Figure 2:**
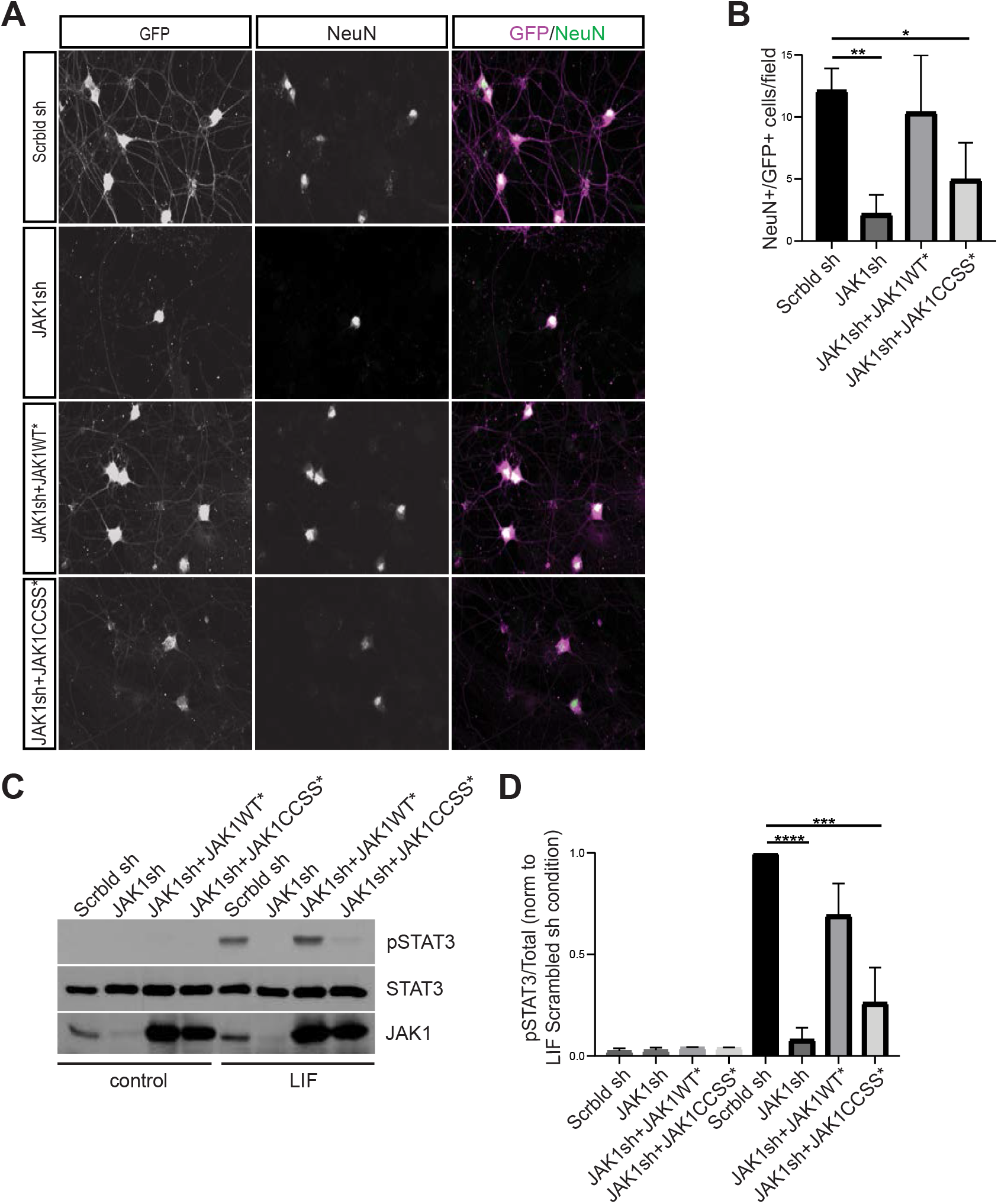
JAK1 palmitoylation is important for DRG neuron survival and signaling in response to neuropoietic cytokines. ***A:*** Images of cultured DRG neurons immunostained with the indicated antibodies 4 days after being plated in the presence of LIF and infected with lentiviruses to express GFP plus a scrambled shRNA (Scrbld sh; top row), GFP plus JAK1 shRNA (JAK1sh; second row), GFP, JAK1shRNA and shRNA-resistant JAK1WT (JAK1WT*; third row) or GFP, JAK1shRNA and shRNA-resistant JAK1-CCSS (JAK1CCSS*; fourth row). ***B*:** Quantified data from *A* confirm greatly reduced numbers of GFP+/NeuN+ neurons after *Jak1* knockdown, which are rescued by shRNA-resistant JAK1WT but not JAK1-CCSS. **: p<0.01, *: p<0.05, one-way ANOVA with Bonferroni *post hoc* test. N=4 determinations per condition ***C:*** Western blots of DRG neurons cultured in the continued presence of NGF to ensure survival, infected with the indicated lentiviruses and subsequently stimulated with LIF. ***D:*** Quantified data from *C* confirm robust LIF-induced STAT3 phosphorylation, which is prevented by *Jak1* knockdown and rescued by JAK1WT* but not JAK1-CCSS*. ****: p<0.0001, ***: p<0.001, oneway ANOVA with Bonferroni *post hoc* test. N=4-5 per condition.

We therefore asked whether LIF-dependent DRG neuron survival requires JAK1 palmitoylation. Consistent with this notion, a second lentivirus expressing an shRNA-resistant form of HA-JAK1WT (JAK1WT*) rescued DRG neuron survival close to the level seen in the presence of the control lentivirus, but a lentivirus expressing palmitoyl-mutant JAK1 (JAK1CCSS*) only minimally rescued neuronal survival (Fig 2A, B). Western blotting of sister cultures trophically supported by Nerve Growth Factor (NGF, which prevents neuronal loss in the absence of JAK1) confirmed similar expression of virally expressed HA-JAK1WT* and HA-JAK1CCSS* (Fig S1A). These findings suggest that JAK1 palmitoylation is important for LIF-induced DRG neuron survival.

The extensive death of *Jak1* ‘knockdown’ and ‘HA-JAK1CCSS*’ neurons in LIF-supported cultures (Fig 2A, B) complicates attempts to define the underlying molecular role(s) of palmitoyl-JAK1. However, when *Jak1* KO neurons from newborn mice are cultured in the presence of NGF they survive equally as well as their wild type counterparts, given their initially lower numbers (Rodig et al. 1998). We therefore used NGF-supported DRG cultures to assess the ability of LIF to trigger phosphorylation of JAK1’s best known substrate, STAT3. Strikingly, both LIF-induced phosphorylation of STAT3 at Y705, the site primarily phosphorylated by JAK, was almost eliminated in *Jak1* knockdown cultures (Fig 2C, D). This result suggested an essential requirement for endogenous JAK1 in LIF-induced STAT3 phosphorylation. LIF-induced STAT3 phosphorylation was fully rescued by HA-JAK1WT*, but was only minimally rescued by HA-JAK1CCSS* (Fig 2C, D). These findings suggest that phosphorylation of endogenous STAT3, and potentially other JAK1 substrates, in response to neuropoietic cytokines requires JAK1 palmitoylation.

### ZDHHC3 and ZDHHC7 are major regulators of JAK1 palmitoylation

We next sought to identify the PAT(s) that controls JAK1 palmitoylation. We therefore screened all 23 mammalian PATs for their ability to palmitoylate co-expressed JAK1 in HEK293T cells. We found that three PATs, the related ZDHHC3 and ZDHHC7, and ZDHHC11, increased JAK1 palmitoylation to a markedly greater extent than any other PAT (Fig 3A, B). Of these PATs, *Zdhhc3* and iZdhhc7 are likely expressed at markedly higher levels than *Zdhhc11* in DRG neurons ((Lerch et al. 2012); www.mousebrain.org). We therefore assessed whether ZDHHC3 and ZDHHC7 are important for endogenous neuronal JAK1 palmitoylation. Indeed, palmitoylation of endogenous JAK1 was markedly reduced in ABE fractions from DRG neurons that had been cultured in the presence of NGF and subsequently lentivirally infected with shRNAs that potently knock down *Zdhhc3* and *Zdhhc7* (Fig 3C, D; effectiveness of shRNAs confirmed in Fig S2). In contrast, *Zdhhc3/7* knockdown did not affect palmitoyl-levels of calnexin, a well-known palmitoyl-protein whose palmitoylation is ascribed to other PATs (Lakkaraju et al. 2012)(Fig 3C, E). These finding suggest that ZDHHC3/7 are essential for endogenous JAK1 palmitoylation in neurons.

**Figure 3:**
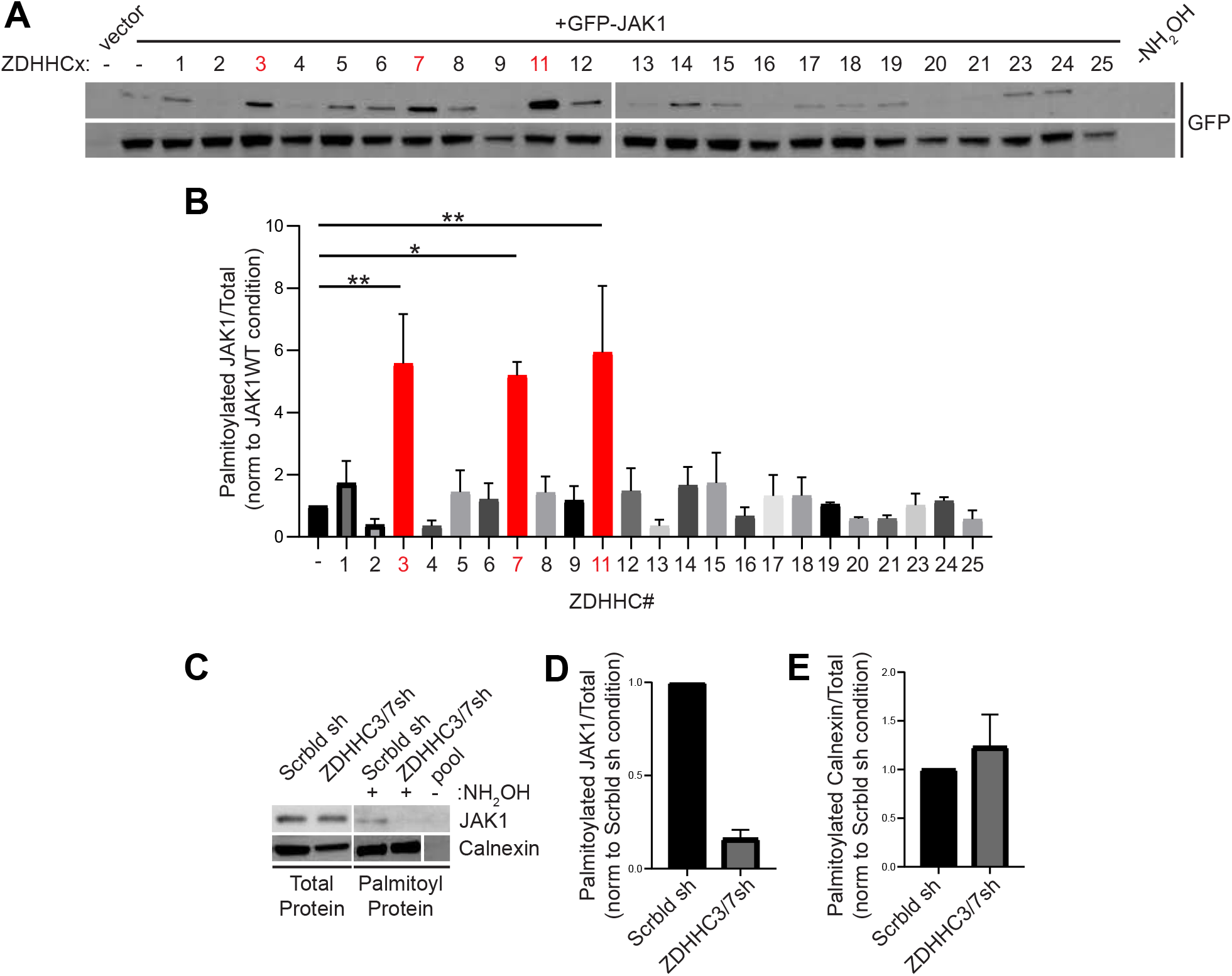
ZDHHC3 and ZDHHC7 dominantly control JAK1 palmitoylation. ***A:*** ABE fractions (top) and total lysates (second panel) from HEK293T cells transfected to express GFP-JAK1 plus the indicated HA-ZDHHC PATs and blotted to detect JAK1. ***B:*** Quantified data from *A* confirm that ZDHHC3, ZDHHC7 and ZDHHC11 most robustly palmitoylated JAK1. **; p<0.01, *;p<0.05 versus JAK1 alone condition, N=3 determinations per condition, one-way ANOVA with Bonferroni *post hoc* test. ***C:*** Total lysates (left panels) and ABE fractions (right panels) from DRG neurons, immunoblotted with the indicated antibodies. ***D, E:*** Quantified data from *C* confirm that ZDHHC3/7 knockdown greatly reduces JAK1 palmitoylation but does not affect calnexin palmitoylation. ***D:*** N=3, Unpaired t-test, p<0.0001. ***E:*** N=3, Unpaired t-test, p=0.5230.

### Palmitoylation is essential for JAK1 kinase activity

Having established the functional importance of JAK1 palmitoylation, we asked how this modification might affect JAK1 at the molecular level. Although palmitoyl-JAK1 is required for STAT3 phosphorylation in neurons, this effect could be due to differential localization of palmitoyl- and depalmitoyl-JAK1. However, in prior work we reported that palmitoylation can also control the intrinsic activity of kinases (Holland et al. 2016). We therefore asked whether palmitoylation might similarly be required for JAK1 to phosphorylate cotransfected STAT3 in HEK293T cells, a non-polarized cell type in which differential subcellular localization is less likely to be a dominant factor, compared to neurons. Phosphorylation of transfected myc-tagged STAT3 (myc-STAT3) at Y705 was undetectable in HEK293T cells, but was markedly upregulated by cotransfected HA-JAK1WT (Fig 4A, B). In contrast, phosphorylation of myc-STAT3 Y705 was also very low in HA-JAK1WT-expressing cells that were treated with 2Br, or in cells cotransfected with HA-JAK1-CCSS (Fig 4A, B). These results suggest that palmitoylation is required for JAK1 to directly phosphorylate STAT3 in cells.

**Figure 4:**
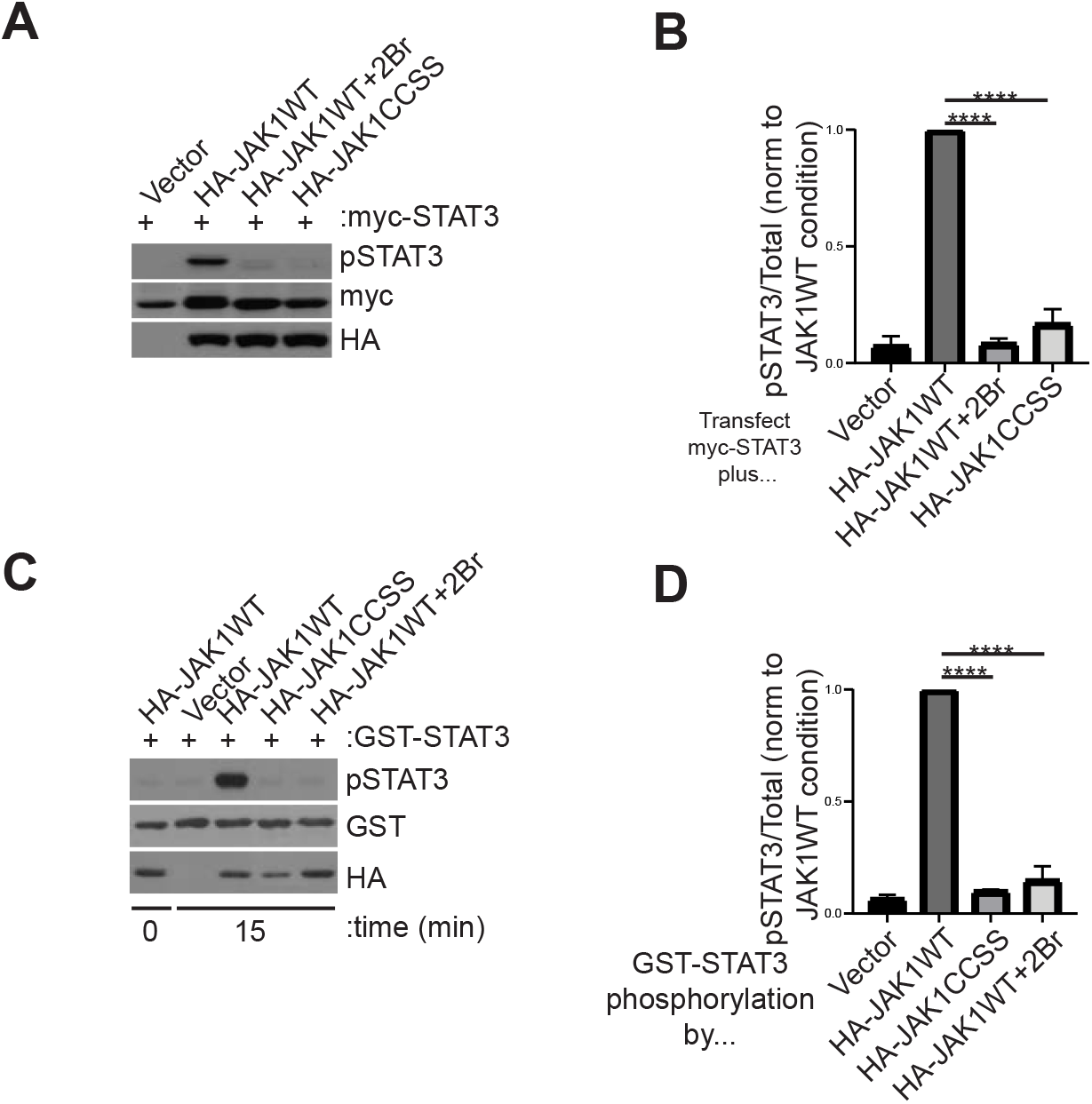
Palmitoylation is essential for JAK1 to phosphorylate STAT3 in cotransfected HEK293T cells and *in vitro*. ***A:*** Western blots of lysates from HEK293T cells transfected with the indicated constructs and treated with or without 2Br prior to lysis. ***B:*** Quantified data from *A* confirm that cotransfection of JAK1 greatly increases STAT3 phosphorylation, an effect prevented by 2Br treatment or CCSS mutation. ****:p<0.0001, one-way ANOVA with Bonferroni *post hoc* test. N=6-11 per condition. ***C:*** Western blots of samples from *in vitro* kinase assays following immunoprecipitation of lysates transfected to express the indicated JAK1 constructs and subsequent incubation with Mg-ATP plus GST-STAT3. Reactions were stopped at the indicated times. ***D:*** Quantified data from *C* confirm robust phosphorylation of GST-STAT3 by JAK1WT, which is prevented by JAK1 CCSS mutation and by 2Br pretreatment. ****:p<0.0001, one-way ANOVA with Bonferroni *post hoc* test. N=3 per condition

Although striking, the palmitoylation-dependent phosphorylation of myc-STAT3 by coexpressed JAK1 (Fig 4A, B) could still potentially be explained by a subcellular localization effect, rather than by a direct requirement of palmitoylation for JAK1 kinase activity. To distinguish between these possibilities, we therefore assessed JAK1’s ability to phosphorylate purified STAT3 in an *in vitro* assay. HA-JAK1WT immunoprecipitated from transfected HEK293T cells robustly phosphorylated GST-tagged STAT3 (GST-STAT3) *in vitro.* In contrast, phosphorylation of GST-STAT3 by HA-JAK1-CCSS, or by HA-JAK1WT isolated from cells pretreated with 2Br, was almost undetectable (Fig 4C, D). These findings suggest that the palmitoylation-dependence of STAT3 phosphorylation by JAK1 is more likely due to effects on JAK1’s catalytic activity rather than its intracellular localization.

We therefore sought to gain more insight into why palmitoylation is necessary for JAK1’s ability to phosphorylate STAT3. To address this question, we used a phospho-specific antibody to assess how palmitoylation affects phosphorylation of tyrosine sites in the activation loop of JAK1’s kinase domain that are critical for JAK1 kinase activity (Y1034, Y1035)(Liu et al. 1997). HA-JAK1WT was robustly phosphorylated at Y1034/Y1035, but phosphorylation of these sites on HA-JAK1CCSS, or on HA-JAK1WT isolated from 2Br-treated cells, was far lower (Fig 5A, B). These results suggest that palmitoylation is essential for JAK1 phosphorylation at Y1034/Y1035.

**Figure 5:**
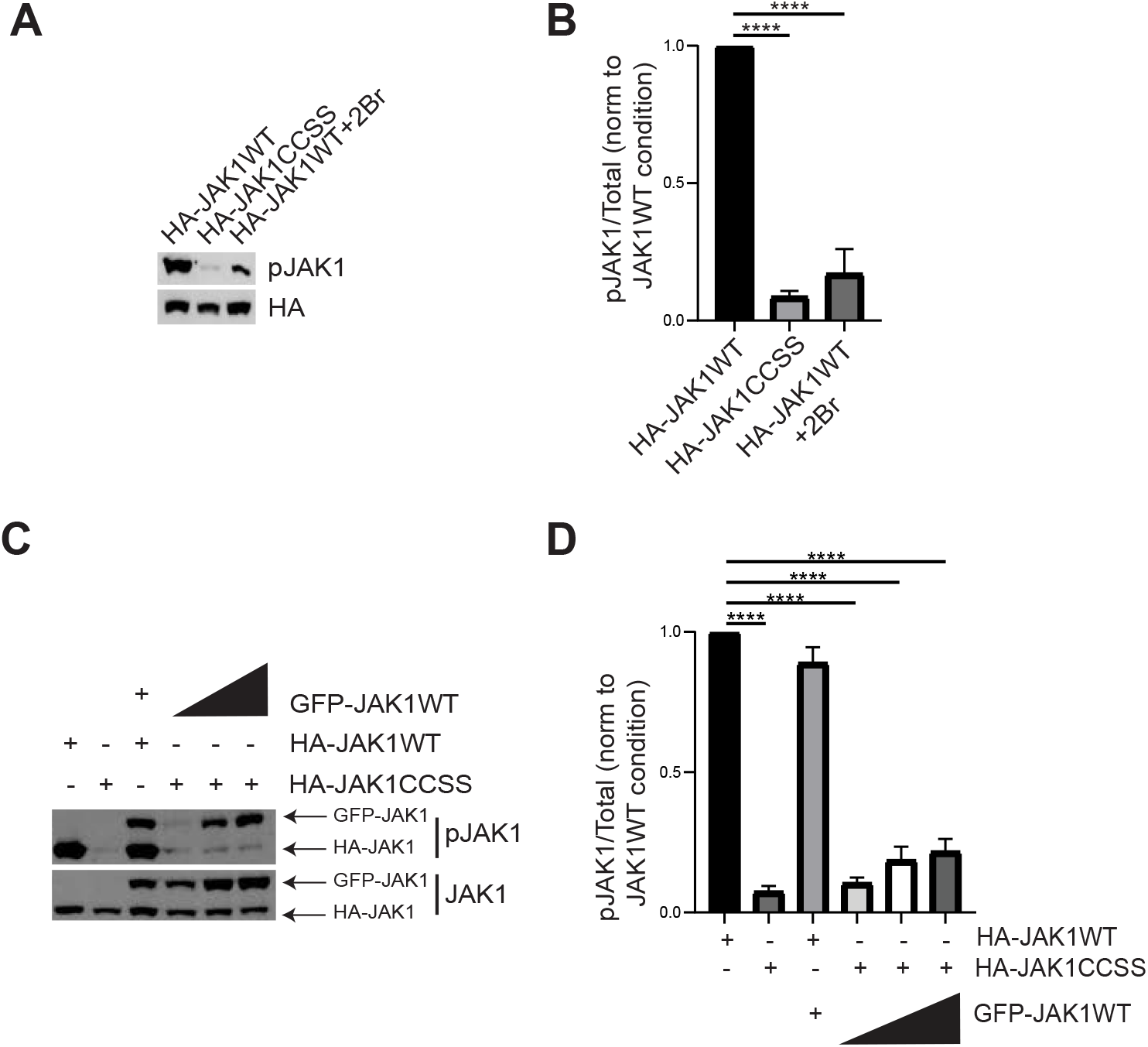
Palmitoylation is required for transphosphorylation of activatory sites on JAK1. ***A:*** Western blots of HEK293T cell lysates transfected to express the indicated JAK1 constructs and treated with or without 2Br. ***B:*** Quantified data from *A* confirm that CCSS mutation and 2Br treatment both greatly reduce JAK1 phosphorylation at Y1034/Y1035. ****:p<0.0001, one-way ANOVA with Bonferroni *post hoc* test. N=3-7 per condition. ***C***: Western blots from HEK293T cells transfected to express the indicated HA-tagged JAK1 constructs with or without GFP-tagged JAK1WT (GFP-JAK1WT). ***D:*** Quantified data from *C* confirm that HA-JAK1 CCSS mutation prevents phosphorylation by GFP-JAK1WT, even when increased amounts of GFP-JAK1 are expressed. ****: p<0.0001, one-way ANOVA with Bonferroni *post hoc* test. N=3-4 per condition.

The prevailing model is that JAK1 becomes activated following transphosphorylation of Y1034/Y1035 within a JAK dimer (Feng et al. 1997; Leonard and O’Shea 1998). Even though JAK1-CCSS is only weakly phosphorylated at these sites when expressed alone, we asked whether this mutant could still be phosphorylated at Y1034/Y1035 by JAK1WT i.e. whether JAK1-CCSS is still potentially ‘activatable’. To address this question, we expressed HA-JAK1WT or HA-JAK1-CCSS with GFP-tagged JAK1WT (GFP-JAK1WT). In lysates from cells expressing HA-JAK1WT and GFP-JAK1WT, we observed strong phospho-JAK1 Y1034/Y1035 signals corresponding to the predicted molecular weights of both these kinases (Fig 5C, D). Consistent with prior experiments, Y1034/Y1035 phosphorylation of HA-JAK1-CCSS expressed alone was very weak and was only minimally increased by co-expressed GFP-JAK1WT (Fig 5C, D). We also observed reduced Y1034/Y1035 phosphorylation of GFP-JAK1WT when coexpressed with HA-JAK1-CCSS, compared to when co-expressed with HA-JAK1WT (Fig 5C). We hypothesized that this reduced phosphorylation might be due to the formation of nonproductive GFP-JAK1WT/HA-JAK1-CCSS dimers in which the reduced activity of HA-JAK1-CCSS precluded transphosphorylation of GFP-JAK1WT at Y1034/Y1035. Consistent with this explanation, we found that expressing higher relative amounts of GFP-JAK1WT:HA-JAK1-CCSS slightly increased phosphorylation of HA-JAK1-CCSS, but more effectively ‘rescued’ the reduced GFP-JAK1WT phosphorylation (Fig 5C, D). These findings suggest that palmitoylation is critical for transphosphorylation of JAK1’s activatory sites, providing a likely explanation for why loss of palmitoylation so markedly reduces JAK1 phosphorylation of exogenous substrates.

## Discussion

Although JAK1 was previously identified as a likely palmitoylated kinase, the importance of this modification for JAK1-dependent signaling and neuronal development has not been previously addressed. Here we report that palmitoylation by the related PATs ZDHHC3 and ZDHHC7 is essential for LIF-dependent signaling and survival of DRG sensory neurons.

Our findings provide broader insights into how JAK signaling controls development of DRG neurons themselves. DRG neuron survival in the presence of neuropoietic cytokines critically requires Gp130-JAK signaling (Nakashima et al. 1999; Rodig et al. 1998). This mechanism of DRG pro-survival signaling is less well studied than that triggered by neurotrophins such as NGF (Harrington and Ginty 2013). However, Gp130/JAK-dependent signaling is physiologically critical for DRG neuron survival *in vivo,* given the reduced number of DRG neurons found in, or isolated from, conventional *Gp130* or *Jak1* KO mice (Nakashima et al. 1999; Rodig et al. 1998). Importantly, the time window during which Gp130/JAK signaling is likely critical *in vivo (*approx. E14-E17 for mice; (Nakashima et al. 1999)) is highly consistent with our findings from cultured neurons, which focus on neurons from E16 rats for their initial four days in culture. It is of note that the *in vivo* time window for this requirement is brief, because perinatal knockout of either *Gp130* or *Jak1* with Nav1.8-Cre, which is active perinatally, has little effect on DRG neuron survival (Andratsch et al. 2009; Oetjen et al. 2017). This result suggests that, just as DRG neurons expressing the NGF receptor *TrkA* become NGF-independent during later development (Rich et al. 1984), the neuropoietic cytokine-dependent pool of DRG neurons rapidly becomes cytokine-independent for survival and/or that other trophic factors can substitute for neuropoietic cytokines at these later timepoints.

It is also interesting that ZDHHC3/7 are the dominant PATs controlling JAK1 palmitoylation. These PATs can palmitoylate an array of substrates (Hayashi et al. 2005; Hayashi et al. 2009; Keller et al. 2004; Kilpatrick et al. 2016). What is therefore surprising is not, perhaps, that ZDHHC3/7 can palmitoylate JAK1, but that other PATs cannot. Notably, both the axonal protein GAP-43 and the GABA receptor gamma2 subunit have been identified as ZDHHC3/7 substrates *in vivo* (Kilpatrick et al. 2016). Both these proteins contain a dicysteine palmitoyl-motif that resembles that of JAK1, suggesting that ZDHHC3/7 effectively palmitoylates motifs of this type. It is also intriguing that *Jak1* KO, *Gp130 (Il6st)* KO and *Zdhhc3/7* DKO mice all die perinatally (Kilpatrick et al. 2016; Nakashima et al. 1999; Rodig et al. 1998), with those that have that been examined thus far exhibiting a marked loss of DRG sensory neurons. There are potentially many reasons for the similarity of these findings, but it is an intriguing possibility that the presence of all these proteins in a common palmitoylation-dependent pathway accounts for this phenotypic overlap.

In seeking to define the importance of palmitoyl-JAK1 signaling, we focused on STAT3, the best described JAK substrate, which (i) mediates a wide variety of ‘downstream’ JAK-dependent events and (ii) causes serious deficits, including death of neuropoietic cytokine-dependent sensory neurons, when knocked out embryonically (Alonzi et al. 2001). Consistent with STAT3 being a key ‘downstream’ palmitoyl-JAK1 substrate, JAK1WT rescues LIF-induced STAT3 phosphorylation and LIF-dependent DRG neuron survival to a very similar extent (Fig 2). However, it remains possible that other substrates also contribute to palmitoyl-JAK1-dependent DRG neuron survival.

Given the likely importance of STAT3 phosphorylation for palmitoyl-JAK1 dependent effects, another key finding of our study is the striking palmitoylation-dependence of JAK1’s kinase activity. JAK1 thus joins a short list of kinases, including Casein Kinase 1 gamma, Fyn, Lck, LIMK1 and DLK, whose function in cells critically requires this modification (Davidson et al. 2005; Montersino and Thomas 2015). Notably, these palmitoyl-kinases fall into two distinct groups. For Fyn, Lck, LIMK1 and CK1gamma, palmitoylation is important for cellular function, but not essential for intrinsic catalytic activity assayed *in vitro* and thus appears to act predominantly as a determinant of kinase localization (Davidson et al. 2005; Montersino and Thomas 2015). In contrast, palmitoyl-mutant forms of DLK and JAK1 not only signal ineffectively in cells but also show vastly reduced kinase activity *in vitro* (Fig 4 and (Holland et al. 2016)). Thus, palmitoylation not only governs localization of these kinases, but also directly controls their enzymatic activity. Why might palmitoylation exert this additional effect on DLK and JAK1? In this regard it is of note that DLK and JAK1 are palmitoylated within the core of their protein sequences, while Fyn, Lck, LIMK1 and CK1gamma are all palmitoylated close to their N-or C-termini. In our prior studies on DLK, we suggested that palmitate addition to DLK’s ‘core’ might have greater ability to alter protein-protein interactions and/or domain structure and hence more directly affect intrinsic catalytic activity (Holland et al. 2016; Montersino and Thomas 2015). However, no reagents (such as phospho-specific antibodies) were available to ‘mark’ DLK’s activation state and further test this hypothesis. Such antibodies are available for JAK1, and reveal that palmitoylation is critical for JAK1 transphosphorylation at Y1034/Y1035, key sites in the activation loop of JAK1’s kinase domain (Liu et al. 1997). We speculate that palmitoylation thus alters JAK1 structure to render these activatory sites more accessible, and/or to increase the intrinsic enzymatic activity of the JAK1 kinase domain, thereby coupling palmitoylation to JAK1’s catalytic activity.

While we have focused our study on neurodevelopmental roles of JAK1 palmitoylation, our findings may also increase understanding of JAK1 function in pathological conditions. In particular, JAK1 expression in developed DRGs is particularly high in pruritogenic neurons that are implicated in chronic itch, and perinatal JAK1 deletion in DRG neurons markedly reduces chronic itch without broadly affecting neuronal function (Oetjen et al. 2017). Consistent with a critical role of JAK1 in mediating chronic itch, pharmacological JAK inhibition reduces chronic itch *in vivo* (Oetjen et al. 2017). However, many JAK inhibitors do not distinguish between JAK1 (which is highly palmitoylated) and other JAK family members (which are likely palmitoylated only at low levels, if at all, but which play key roles in other cell types). It is an intriguing possibility that pharmacologically targeting JAK1 palmitoylation, rather than broadly inhibiting JAK kinase activity, could provide a means to alleviate chronic itch without affecting other physiological roles of JAK signaling.

Our findings that palmitoylation of JAK1 is essential for DRG neuron survival provides new insights into how signaling pathways are controlled during neurodevelopment. The tight coupling of JAK1’s palmitoylation and kinase activity suggest that JAK1 is only active when membrane-associated. Given the location of JAK1’s palmitoylation sites in a flexible linker region that is altered by *JAK1* oncogenic mutation (Toms et al. 2013), it will be of great interest to assess how and whether JAK1 palmitoylation levels may be altered in genetic and idiopathic disease.

## Materials and Methods

### Antibodies

The following antibodies, from the indicated sources, were used in this study: rabbit anti-phospho-STAT3 (#9145), rabbit anti-STAT3 (#8768), rabbit anti phospho-JAK1 (#74129), rabbit anti-myc (#9145), all from Cell Signaling Technology, Danvers, MA; mouse anti-myc 9E10 (UPenn Cell Center); rabbit anti-GFP #A11122 (Invitrogen); mouse anti-JAK1 #610231 (BD Transduction Labs), mouse-anti NeuN #MAB377 (Millipore Sigma); mouse anti HA11 #901514 (Biolegend), Calnexin #10286 (Abcam). Unless otherwise noted, all chemicals were from ThermoFisher Biosciences and were of the highest reagent grade available.

### HEK293T Cell Culture and Transfection

HEK293T cells were cultured in Dulbecco’s Modified Eagle Medium (DMEM, Thermo Fisher Scientific) with 10% fetal bovine serum (FBS), 1% Penicillin-Streptomycin (Thermo Fisher Scientific), and 1x GlutaMAX (Thermo Fisher Scientific). A calcium phosphate method described previously (Thomas et al. 2005) was used for all HEK293T cell transfections.

### DRG Neuron Culture

Dorsal root ganglion (DRG) neurons were dissociated from E16 rat embryos and conventional ‘mass’ cultures were prepared as previously described (Holland et al. 2016), except that in some experiments 25 μg/ml LIF (Alomone Labs) was used as trophic support rather than NGF. Neurons were re-fed the day after plating (1 day *in vitro;* DIV1) with Neurobasal medium containing NGF or LIF plus B27 and Fluorodeoxyuridine to inhibit growth of mitotic cells.

Neurons were infected with lentiviruses on DIV3 and used for experiments on DIV6. NGF-containing cultures were stimulated wit 1 ng/ml (final concentration) LIF.

### Lentiviral Vectors and Preparation

A *Jak1* shRNA (5’-gcctgagagtggaggtaac-3’) was identified bioinformatically and synthesized with a neighboring shRNA promoter by Genewiz. The resultant cassette was subcloned into lentiviral vector FEGW (Holland et al., 2016). Human JAK1 cDNA was obtained from DNASU and subcloned into HA-FEW or GFP-FEW vectors using primers with *XhoI* and *NotI* restriction sites. Human *JAK1* cDNA was made shRNA-resistant by subcloning a gene synthesized *BsrG/-NotI* fragment (Genewiz) in which shRNA recognition sequence was mutated while maintaining protein coding sequence. CCSS mutation was introduced by Quickchange mutagenesis. Human STAT3 cDNA was also obtained from DNASU and was subcloned into myc-FEW and pCIS-GST vectors, the latter vector allowing production of GST-tagged STAT3 fusion protein in mammalian cells. Mouse *Zdhhc* cDNAs were a kind gift of Dr. Masaki Fukata and were subcloned into HA-FEW vector as described (Holland et al. 2016). ShRNA sequences against rat *Zdhhc3* (5’-GAGACATTGAACGGAAACCAGAATACCTC-3’) and *Zdhhc7* (5’-ATGACATGGCTTCTGGTCGTCTATGCAGA-3’) were purchased from Origene and subcloned, together with their neighboring U6 cassette, into FEGW. VSV-G pseudotyped lentiviruses were made in HEK293T cells as previously described (Thomas *et al.,* 2012).

### Acyl Biotin Exchange Assay (ABE)

Palmitoylation was measured by acyl biotin exchange assay as described previously (Thomas et al. 2012). Briefly, HEK293T cells or DRG neurons were lysed in ABE lysis buffer (50 mM HEPES, pH 7, 2% [w/v] sodium dodecyl sulfate (SDS), 1 mM EDTA plus protease inhibitors (PIC)) and 20 mM thiol-reactive methyl-methane thiosulfonate (MMTS), sonicated, and incubated at 50 degrees Celsius for 20 minutes. Protein was precipitated by addition of chilled acetone (80% final [v/v]), and MMTS was removed by sequential washes with 80% [v/v] acetone. Pellets were resuspended in 4% SDS buffer (4% [w/v] SDS, 50 mM Tris pH 7.5, 5 mM EDTA plus PIC) and a fraction was removed as an ‘Input’ sample. Inputs were taken out in dilution buffer (50 mM HEPES, 1% [v/v] triton X-100, 1 mM EDTA, 1 mM EGTA) plus PIC, 150 mM NaCl and 5x sample buffer with β-mercaptoethanol (BME) (Millipore-Sigma). Samples were split in two and incubated for 1 hour rotating in the dark at room temperature in 1xProtease Inhibitor Cocktail (Boehringer), 1 mM HPDP-biotin (Soltec Ventures, Beverly, MA), 0.2% Triton X-100, with either 1M hydroxylamine pH 7.5 or 50 mM Tris pH 7.5. Samples were acetone precipitated to remove excess hydroxylamine and HPDP-biotin and pellets were resuspended in ABE lysis buffer plus PIC and diluted 1:20 in dilution buffer plus 150 mM NaCl and protease inhibitors. Biotinylated proteins were captured by incubation with high capacity neutravidin-conjugated beads (Thermo Fisher Scientific) for 3 hours at 4 degrees Celsius. Beads were washed three times with dilution buffer containing plus 0.5 M NaCl, and twice with dilution buffer alone. Protease inhibitors (4 μg/ml Leupeptin and 1 mM Benzamidine) were added to all washes. Proteins were eluted from beads by addition of 1% (v/v) BME, 0.2% [w/v] SDS, 250 mM NaCl in dilution buffer and incubated for 10 minutes at 37 degrees Celsius. Supernatants were removed and denatured by adding one-fifth volume of 5x SDS sample buffer. Samples were boiled and subjected to SDS-PAGE and western blotting.

### Purification of GST-STAT3

HEK293T cells were transfected with pCIS-GST-STAT3 vector and cells were lysed 24 h later in immunoprecipitation buffer (IPB: 1x phosphate buffered saline [PBS], 1% [w/v] Triton X-100, 50 mM NaF, 5 mM Na_4_P_2_O_7_, 1 mM Na_3_VO_4_, 1 mM EDTA, and 1 mM EGTA) plus 1x PIC. Lysates were centrifuged at 13,000 x *g* and supernatants incubated with Glutathione Sepharose beads (pre-equilibrated with IPB) for 90 min at 4 degrees C on a rotating platform. Beads were then washed 4 times with IPB plus 0.25 M NaCl, twice with IPB and once with 50 mM Tris pH 7.5. All wash buffers contained protease inhibitors (4 μg/ml Leupeptin and 1 mM Benzamidine). GST-tagged STAT3 was eluted by addition of 20 mM Glutathione pH8.0 plus 250 mM NaCl and was dialyzed extensively against 50 mM Tris pH 7.5 containing 50%(v/v) glycerol. GST-STAT3 was stored unfrozen at −20 degrees C. The preparation was judged to be >80% pure and was used for subsequent *in vitro* kinase assays.

### Immunoprecipitation-Kinase Activity Assay

Lysates from HEK293T cells transfected to express HA-tagged JAK1WT or -CCSS were treated with or without 20 μM 2Br and lysed in IPB. Lysates were spun at 13000 rpm for 10 minutes at 4 degrees Celsius. Supernatant was then filtered through a SpinX column. A fraction of each sample was denatured in SDS sample buffer for use as ‘Inputs’. 200 microliters of the remaining lysate was incubated with 5 μl of Protein G Sepharose beads (settled volume) that had been pre-coupled with 1 μl of anti-HA ascites. This antibody-bead complex was incubated for 90 minutes at 4 degrees C while rotating. Beads were washed twice with IPB containing and 0.5 M NaCl, twice with IPB alone and once with 1x Kinase assay buffer (10 mM Tris pH 7.5, 0.2 mM EDTA, and 0.1% TritonX-100). Excess supernatant was removed. Samples were then mixed with the Kinase Activity Assay components, with the final reaction containing of beads, 1 μM GST-STAT3, 0.1 mM ATP, 10 mM MgCl_2_ in 1x Kinase assay buffer. A fraction of the reaction mix was immediately removed as a t=0 sample and reactions were then allowed to proceed at room temperature with regular flick-mixing. Fractions of the reaction were removed and stopped by dilution in 1x SDS sample buffer, boiled for 5 minutes and subjected to SDS-PAGE and western blotting.

### Immunostaining

For immunocytochemical readouts, dissociated DRG neurons cultured on coverslips were rinsed once with 1x Recording buffer (25 mM HEPES pH7.4, 120 mM NaCl, 5 mM KCl, 2 mM CaCl_2_, 1 mM MgCl_2_, 30 mM Glucose) and fixed in 4% paraformaldehyde (PFA)/sucrose for 10 min at room temperature. Samples were permeabilized in PBS containing 0.25% (w/v) Triton-X-100 for 10 min at 4°C, blocked with PBS containing 10% (v/v) Normal Goat Serum (SouthernBiotech, 0060-01) for 1 h and incubated in primary antibodies overnight at 4°C in blocking solution. After 3 washes with PBS, cells were incubated for 1 h at room temperature with AlexaDye-conjugated fluorescent secondary antibodies diluted in blocking solution, prior to 3 final PBS washes and mounting in FluorSave reagent (Millipore Sigma).

### Image Acquisition

Images were acquired using a Nikon C2 confocal microscope (20x, 0.75NA objective) and NIS Elements software. Files were exported and quantified using Fiji/ImageJ. Images shown are maximum intensity projections of confocal Z-stacks.

## Supporting information

Combined Supplementary Data file for Hernandez et al.

## Acknowledgments

We thank Drs. Eric Witze (University of Pennsylvania) for comments on the manuscript and Masaki Fukata (National Institute of Physiological Sciences, Okazaki, Japan for PAT cDNAs). Supported by grants from NIH (R01NS094402) and Shriners Hospitals for Children (85600PHI) to G.M.T.

