## Supplementary material for "JAK1 palmitoylation by ZDHHC3/7 is Essential for Neuropoietic Cytokine Signaling and DRG Neuron Survival": Combined Supplementary Data file for Hernandez et al.

**Supplementary Figure Legends:**

**Figure S1. Confirmation of JAK1 knockdown and rescue and requirement of palmitoylation for CNTF-induced JAK1 signaling.** ***A:*** Western blots of DRG neuron lysates infected with the indicated lentiviruses. JAK1 shRNA lentivirus greatly reduces JAK1 protein levels, which are rescued by shRNA-resistant JAK1WT or –CCSS (JAK1WT*, JAK1CCSS*). JAK1WT* and JAK1CCSS* express at broadly similar levels.

**Figure S2. Confirmation of effective knockdown of cognate targets by *Zdhhc3* and *Zdhhc7* shRNAs.** ***A:*** HEK293T cells were cotransfected to express HA-tagged rat ZDHHC3 plus either a scrambled shRNA or shRNA against rat *Zdhhc3.* The shRNA greatly reduces ZDHHC3 protein levels and was used for subsequent experiments in neurons. ***B:*** As *A*, except that HEK293T cells were cotransfected to express HA-tagged ZDHHC7 plus either a scrambled shRNA or one of four shRNAs against rat *Zdhhc7*. The second of these shRNAs (ZDHHC7shB) most effectively reduced ZDHHC7 protein levels and was used for subsequent experiments in neurons.

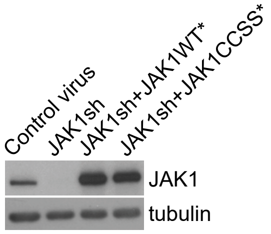
Control virus
JAK1sh
JAK1sh+JAK1WT*
JAK1sh+JAK1CCSS*
JAK1
tubulin

**A**

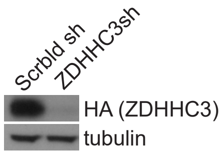
Scrbld sh
ZDHHC3sh
HA (ZDHHC3)
tubulin

**B**

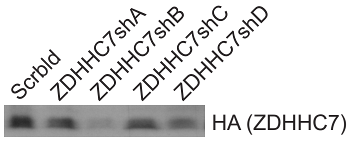
Scrbld
ZDHHC7shA
ZDHHC7shB
ZDHHC7shC
ZDHHC7shD
HA (ZDHHC7)
